# Rab27b promotes lysosomal function and alpha-synuclein clearance in neurons

**DOI:** 10.1101/2024.06.20.599785

**Authors:** Kasandra Scholz, Rudradip Pattanayak, Ekkatine Roschonporn, Frank Sanders Pair, Amber Nobles, Talene A. Yacoubian

## Abstract

Alpha-synuclein (αsyn) is the key pathogenic protein implicated in synucleinopathies including Parkinson’s Disease (PD) and Dementia with Lewy Bodies (DLB). In these diseases, αsyn is thought to spread between cells where it accumulates and induces pathology; however, mechanisms that drive its propagation or aggregation are poorly understood. We have previously reported that the small GTPase Rab27b is elevated in human PD and DLB and that it can mediate the autophagic clearance and toxicity of αsyn in a paracrine αsyn cell culture neuronal model. Here, we expanded our previous work and further characterized a role for Rab27b in neuronal lysosomal processing and αsyn clearance. We found that Rab27b KD in this αsyn inducible neuronal model resulted in lysosomal dysfunction and increased αsyn levels in lysosomes. Similar lysosomal proteolytic defects and enzymatic dysfunction were observed in both primary neuronal cultures and brain lysates from Rab27b knockout (KO) mice. αSyn aggregation was exacerbated in Rab27b KO neurons upon treatment with αsyn preformed fibrils. We found no changes in lysosomal counts or lysosomal pH in either model, but we did identify defects in acidic vesicle trafficking in Rab27b KO primary neurons which may drive lysosomal dysfunction and promote αsyn aggregation. Rab27b OE enhanced lysosomal activity and reduced insoluble αsyn accumulation. Finally we found elevated Rab27b levels in human postmortem incidental Lewy Body Disease (iLBD) subjects relative to healthy controls. These data suggest a role for Rab27b in neuronal lysosomal activity and identify it as a potential therapeutic target in synucleinopathies.

## INTRODUCTION

As the global population ages, synucleinopathies including Parkinson’s Disease (PD) and Dementia with Lewy Bodies (DLB) are expected to increase dramatically in prevalence, exacerbating the existing social and economic burdens associated with patient care [1–3]. These diseases are pathologically characterized by intracellular proteinaceous inclusions comprised of alpha-synuclein (αsyn), which misfolds, spreads between cells, aggregates, and induces pathology [4, 5]. It is thought to spread in a prion-like manner where toxic, oligomeric species contact healthy monomeric αsyn and template its misfolding [6, 7], but the mechanisms governing this spread and accumulation remain poorly understood.

Various biological processes that likely contribute to the spread and aggregation of αsyn include autophagic and endolysosomal pathways. These pathways are active areas of interest in Parkinson’s Disease; both monomeric and misfolded αsyn can be degraded by autophagic processes [8, 9], PD pathology is associated with autophagic-lysosomal dysfunction [5, 10], and upregulating autophagy may have therapeutic potential [11, 12]. In disease states, dysregulated autophagy may impair αsyn clearance and thereby facilitate aggregation and cell death [13–15]. Further, αsyn itself may damage autophagic capacity, resulting in a positive feedback loop that can exacerbate pathology [16, 17]. Within these pathways, lysosomes are key structures of interest, and lysosomal impairment may be a major contributor to PD pathogenesis [18–20]. Several genetic lysosomal defects result in or increase the risk of PD, including mutations in the lysosomal ATPase *ATP13A2* and the lysosomal enzyme *GBA1* [21, 22]. Autophagic-lysosomal pathways, which are regulated in part by members of the Rab family of GTPases, may therefore play a critical role in disease progression.

Rab GTPases are a family of over 60 proteins that regulate a variety of cellular processes including trafficking, endocytic, secretory, and autophagic-lysosomal pathways [23–25]. Several members of this family have been implicated in neurodegeneration and PD, both as direct modulators of αsyn and targets of the PD-associated kinase *LRRK2* [26–29]. Rab27b is a member of this family that has been implicated in multiple cellular processes including exosome secretion and synaptic transmission [30, 31]. We have previously identified Rab27b as a regulator of the release, toxicity, and clearance of αsyn in an *in vitro* inducible human αsyn model [32]. Additionally we found an increase in Rab27b levels in the temporal cortex of human PD and DLB patients relative to age-matched controls [32]. Here we describe a new role for Rab27b in the lysosomal function of neurons and in the lysosomal clearance of αsyn. We evaluated Rab27b’s role in lysosomal function and αsyn pathology using knockdown (KD), knockout (KO), and overexpression (OE) models *in vitro* and *in vivo*. We found that KD or KO of Rab27b impairs lysosomal function and exacerbates αsyn pathology, while OE conversely enhances lysosomal functioning and ameliorates pathology, establishing Rab27b as a regulator of autophagic-lysosomal processing in the context of synucleinopathies.

## RESULTS

### Rab27b KD promotes αsyn accumulation in lysosomes

We previously developed an inducible cell culture model that overexpresses and secretes human αsyn when exposed to doxycycline (doxy) [33]. This model is hereafter referred to as “isyn” for inducible synuclein. We knocked down Rab27b in this model using a puromycin-selected lentiviral shRNA construct and additionally generated a scrambled shRNA non-targeted (nt) control cell line [32]. We have previously shown that KD of Rab27b in these cells exacerbates αsyn pathology and toxicity, with accumulation of Triton X-100 insoluble αsyn [32]. We also found that Rab27b KD increased levels of p62 and LC3B-positive puncta, suggesting a role in autophagic-lysosomal pathways. To examine where in the autophagic-lysosomal pathway Rab27b may act, we first performed colocalization experiments of Rab27b with proteins associated with autophagic-lysosomal pathways. We treated isyn cells with doxy for 96 hours to induce αsyn overexpression before staining for Rab27b in combination with the autophagosomal marker LC3B, autophagic substrate marker p62, and the lysosomal marker LAMP-1. We found that Rab27b partially colocalized with all three autophagic-lysosomal markers (Fig. 1a-d). Additionally, we assessed the interaction of Rab27b with αsyn via proximity ligation assay and found that it interacts with αsyn in induced isyn cells (Fig. 1e). This finding that Rab27b partially colocalizes with several markers in the autophagic-lysosomal pathway suggests that Rab27b may be involved in the trafficking of vesicles as they transition through this pathway.

**Figure 1.**
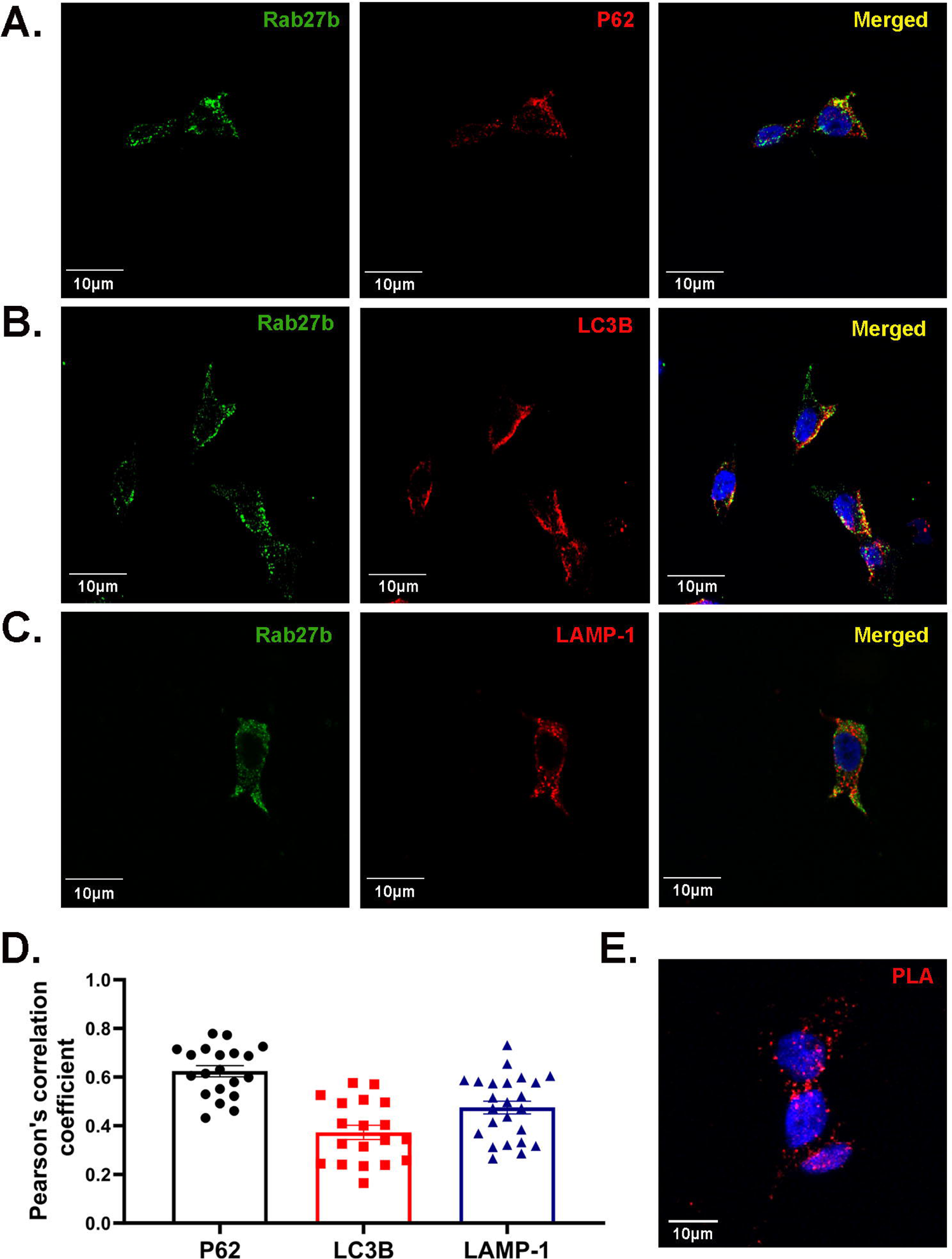
Rab27b partially colocalizes with autophagic-lysosomal markers and interacts with αsyn. A-D: Rab27b (green) partially colocalizes with p62, LC3B, and LAMP-1 (red) in isyn cells induced with doxy for 96 hours. Shown are representative images and quantification of individual cells across three rounds of staining. Error bars denote SEM. B: Stain showing PLA-positive puncta (red) indicating instances of Rab27b interaction with αsyn.

Given Rab27b’s association with lysosomal structures, we next performed lysosomal fractionation to determine if αsyn accumulates within lysosomes upon Rab27b KD in isyn cells. We isolated lysosomes from doxy-induced nt-shRNA control isyn and isyn/Rab27b KD cells via ultracentrifugation and assessed the amount of αsyn found in the lysosomally-enriched fractions via Western blotting. We found that αsyn levels in lysosomal fractions were higher in isyn/Rab27b KD cells compared to control cells (Fig. 2a). As previously reported [32], total lysate levels between control and isyn/Rab27b KD cells were not different. These data indicate that increased αsyn levels within the lysosomal fractions of isyn/Rab27b KD cells could be secondary to defects in αsyn lysosomal degradation.

**Figure 2.**
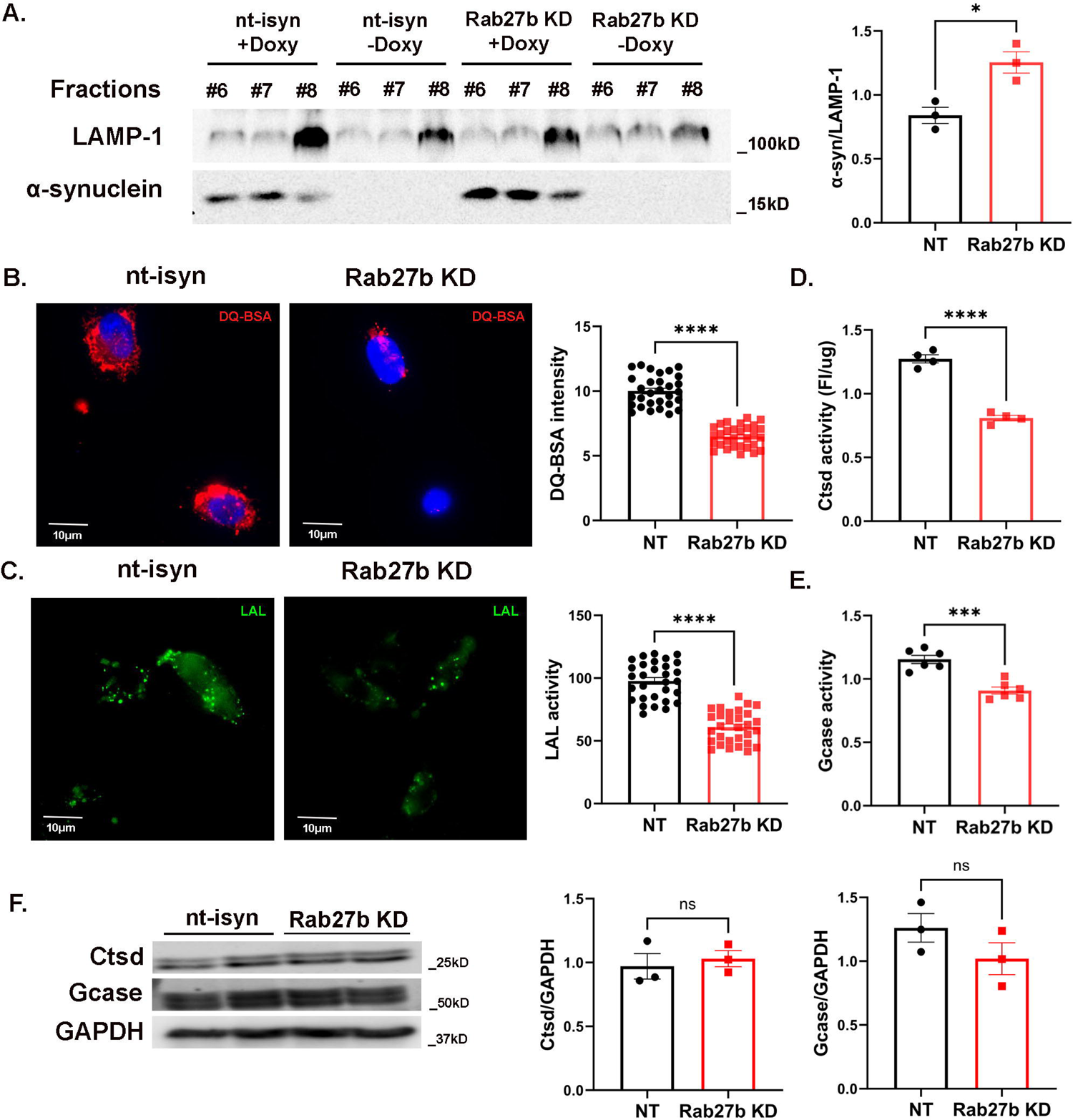
Rab27b KD causes lysosomal impairments in inducible αsyn cells. A: Representative Western blot showing increased αsyn levels in LAMP-1-enriched fractions of isyn/Rab27b KD cells relative to control cells. Fractions 6-8 denote successive fractions. αSyn levels were normalized to LAMP-1 enrichment. n=3 independent rounds. Student’s t-test; *p<0.05. B: Representative images and quantification of DQ-BSA activity in nt-isyn and isyn/Rab27b KD cells. n=30 individual cells across 3 independent rounds. Student’s t-test; ****p<0.0001. C: Representative images and quantification of LAL activity in nt-isyn and isyn/Rab27b KD cells. n=30 individual cells across 3 independent rounds. Student’s t-test; ****p<0.0001. D: Quantification of Ctsd activity in nt-isyn and isyn/Rab27b KD cells. n=4 independent rounds. Student’s t-test; ****p<0.0001. E: Quantification of Gcase activity in nt-isyn and isyn/Rab27b KD cells. n=6 independent rounds. Student’s t-test; ***p<0.001. F: Representative Western blot and quantification of mature Ctsd and Gcase expression in nt-isyn and isyn/Rab27b KD cells. n=3 independent rounds. Student’s t-test. All error bars denote SEM.

### Rab27b is required for lysosomal function in a doxy-inducible αsyn cell line

Given the accumulation of αsyn within lysosomes upon Rab27 KD in isyn cells, we next assessed lysosomal function isyn/Rab27b KD cells. We first measured generalized lysosomal proteolysis by treating cells with an autoquenched DQ-BSA dye, which fluoresces only after exposure to active proteases. We found that isyn/Rab27b KD cells showed 43% reduction in DQ-BSA signal relative to control cells six hours post-treatment (Fig. 2b), indicating generalized lysosomal impairment. We then assessed the activity of several lysosomal enzymes using quenched substrates specific to each enzyme. Isyn/Rab27b KD cells showed a 47% reduction in lysosomal acid lipase (LAL) activity compared to nt-shRNA control cells (Fig. 2c). Similarly, Rab27b KD in isyn cells reduced cathepsin D (Ctsd) and glucocerebrosidase (Gcase) activities compared to nt-shRNA control isyn cells (Fig. 2d-e). We next tested whether reduced lysosomal activity with Rab27b KD was secondary to a reduction in enzyme protein levels and found no changes in the expression levels of Gcase or mature Ctsd protein (Fig. 2f).

To determine if these effects were driven by a reduction in lysosomal content, we then measured the number of lysosomes in each cell line using the lysosome-specific dye Lysotracker Red. We found no differences in the number of Lysotracker-positive vesicles in isyn/Rab27b KD cells compared to nt-shRNA control cells (Fig. 3a). To validate this, we also assessed lysosomal numbers using LAMP-1 immunocytochemistry. LAMP-1-positive vesicles were similar in nt-shRNA control and isyn/Rab27b KD cells (Fig. 3b). These data indicate that the observed impairments in lysosomal function were not due to a reduction in the number of lysosomes.

**Figure 3.**
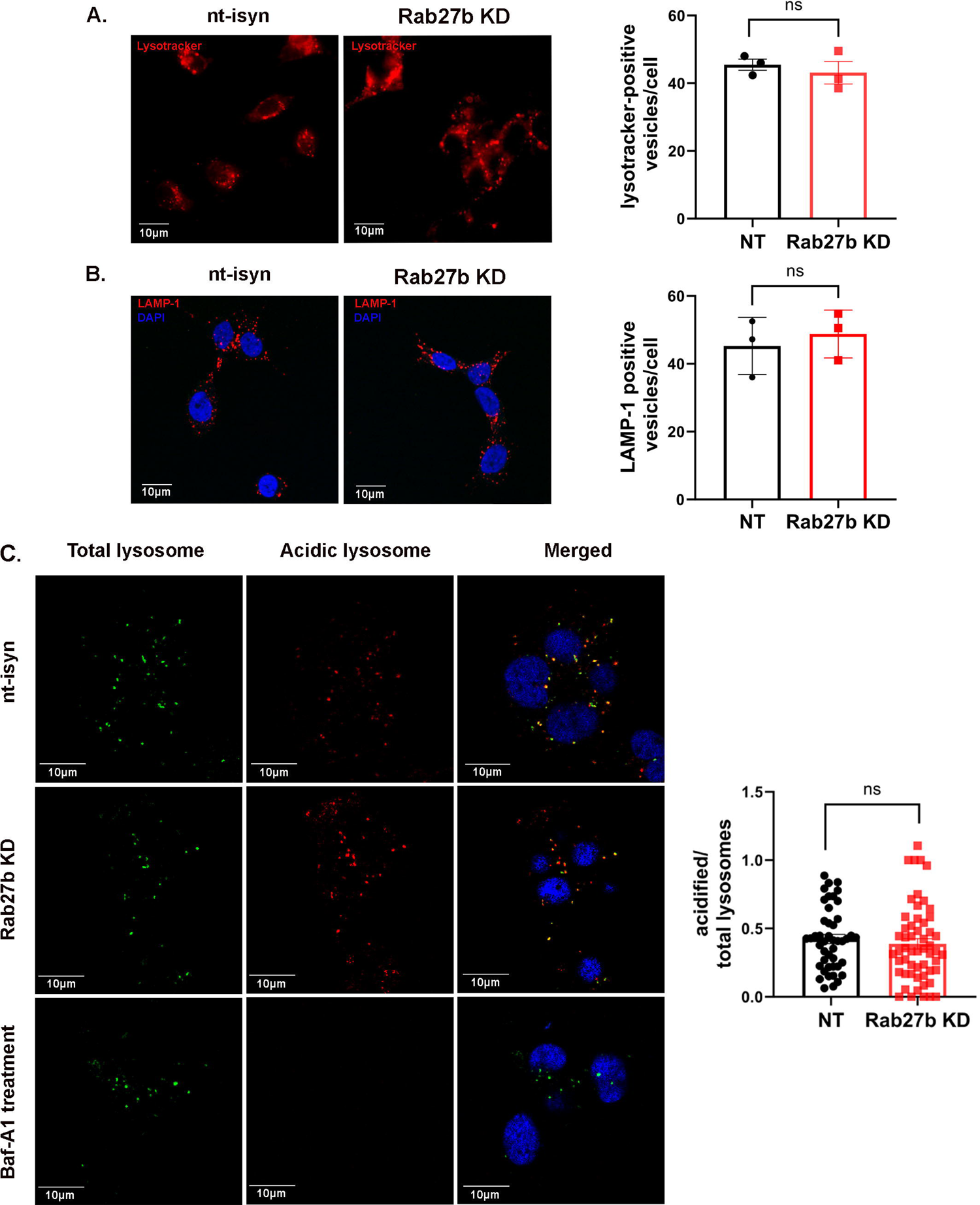
Rab27b KD does not affect lysosomal counts or pH in isyn cells. A: Representative images and quantification of Lysotracker in nt-isyn control and isyn/Rab27b KD cells. n= 3 independent rounds. Student’s t-test. B: Representative images and quantification of LAMP-1-positive puncta in nt-isyn control and isyn/Rab27b KD cells. n=3 independent rounds. Student’s t-test. C: Representative images of total lysosomes (green) and acidified lysosomes (red) in nt-isyn control and isyn/Rab27b KD. As a negative control, some cells were treated with BafA to inhibit lysosomal acidification. n=45-55 cells across 3 independent rounds. Student’s t-test. All error bars denote SEM.

We next assessed whether lysosomal pH was altered upon Rab27b KD in isyn cells. Living isyn/Rab27b KD and control cells were induced with doxy for 96 hours before being stained and quantified for both total and acidified lysosomes. A subset of cells were treated with Bafilomycin A (BafA) to inhibit acidification as a negative control. Isyn/Rab27b KD cells showed a similar proportion of acidified lysosomes relative to the total number compared to nt-shRNA control isyn cells (Fig. 3c). Overall, isyn/Rab27b KD cells exhibited defects in lysosomal enzymatic activity but no changes in lysosomal counts, lysosomal pH, or enzyme expression levels.

### Lysosomal activity is impaired in the brains of Rab27b knockout mice

We next examined lysosomal function in primary neurons from Rab27b knockout (KO) mice. We generated Rab27b single KO mice after crossing Rab27a/Rab27b double KO (DKO) mice obtained from Dr. Seabra [34] with wildtype (WT) C57BL/6J mice. Of note, we previously found that the Rab27a/b DKO mouse carries a spontaneous deletion that includes *Snca*, the gene that encodes αsyn, and *Mmrn1*, the gene that encodes multimerin 1 [35]. This deleted region on chromosome 6 was restored in a subset of progeny after crossing the DKO mice with C57BL/6J mice. Mice heterozygous for Rab27a and Rab27b that lacked the deletion region were selected for subsequent breeding in order to generate Rab27b single KO mice (Fig. 4a). We confirmed that this previously deleted region, including the *Snca* locus, was present in the Rab27b single KO mice (Fig. 4b). Interestingly, in our attempts to generate Rab27a/Rab27b double KO (Rab27 DKO) mice without the deletion, we found that KO of both Rab27a and Rab27b was not compatible with the presence of the *Snca/Mmrn1* region. Only three Rab27a/b DKO mice were born out of a total of 97 pups born from mating Rab27a ^-/-^ Rab27b ^-/-^ αsyn^+/-^ mice to each other, which was much less than the expected 25% (Table 1). Additionally, we attempted three other crosses to generate the Rab27 DKO mouse line without the *Snca/Mmrn1* deletion, yet these crosses also primarily produced fewer than expected Rab27 DKO pups (Table 1).

**Figure 4.**
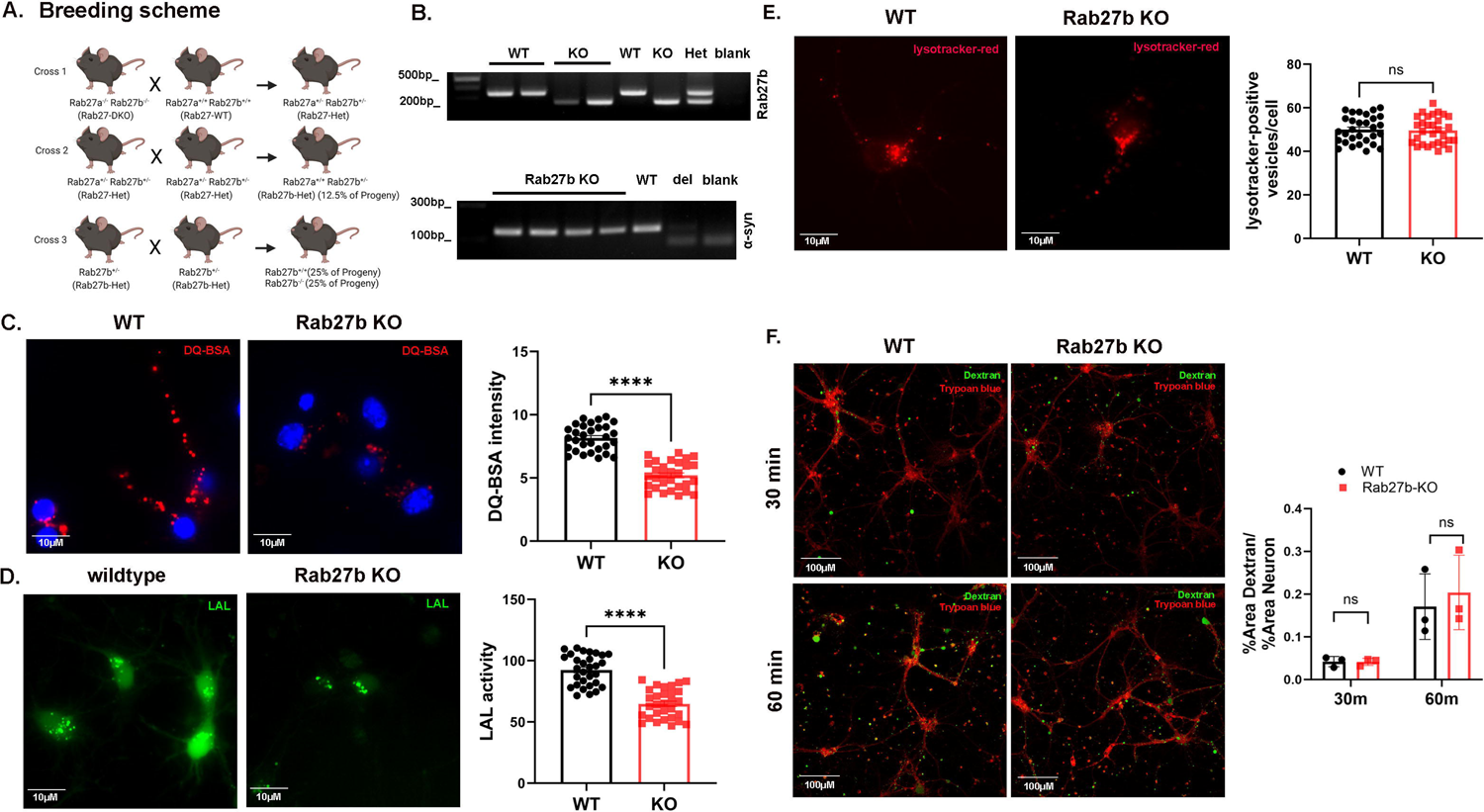
Rab27b KO primary neurons display lysosomal deficits but no changes in lysosomal number or uptake capabilities. A: Schematic depicting Rab27b KO breeding strategy. B: Rab27b KO line genotyping validating that Rab27b KO and littermate WT mice express the *Snca* locus. C: Representative images and quantification of DQ-BSA activity in WT and Rab27b KO neurons. n=30 individual cells across 3 independent rounds. Student’s t-test; ****p<0.0001. D: Representative images and quantification of LAL activity in WT and Rab27b KO neurons. n=30 individual cells across 3 independent rounds. Student’s t-test; ****p<0.0001. E: Representative images and quantification of Lysotracker-positive puncta in WT and Rab27b KO neurons. n=30 individual cells across 3 independent rounds. Student’s t-test. F: Representative images and quantification of 10kDa 488-Dextran uptake in WT and Rab27b KO neurons at 30 and 60 minutes. n=3 independent rounds. Student’s t-test. All error bars denote SEM.

**Table 1.** Crosses to generate Rab27 DKO mice result in lower than expected proportions of DKO mice.

| Breeding cross | Expected proportion for Rab27 A<sup>-/-</sup> Rab27 B<sup>-/-</sup> syn<sup>+/+</sup> | Actual proportion for Rab27 A<sup>-/-</sup> Rab27 B<sup>-/-</sup> syn<sup>+/+</sup> |
| --- | --- | --- |
| A<sup>+/-</sup> B<sup>+/-</sup> syn <sup>+/+</sup> X<br>A<sup>+/-</sup> B <sup>+/-</sup> syn <sup>+/+</sup> | 6.25% | 3.2% (1 of 31 pups) |
| A<sup>-/-</sup> B<sup>-/-</sup> syn <sup>+/+</sup> X<br>A<sup>-/-</sup> B <sup>+/-</sup> syn <sup>+/+</sup> | 50% | 0% (0 of 14 pups) |
| A<sup>+/-</sup> B<sup>+/-</sup> syn <sup>+/-</sup> X<br>A<sup>+/-</sup> B <sup>+/-</sup> syn <sup>+/+</sup> | 3.1% | 8.3% (3 of 36 pups) |
| A<sup>-/-</sup> B<sup>-/-</sup> syn <sup>+/-</sup> X<br>A<sup>-/-</sup> B<sup>-/-</sup> syn <sup>+/-</sup> | 25% | 3.1% (3 of 97 pups) |

Once we established the single Rab27b KO mouse line, we first examined lysosomal function in primary hippocampal neurons at *day in vitro* (DIV) 8. Lysosomal proteolytic activity as assessed by DQ-BSA cleavage was reduced by 45% in neurons from Rab27b KO mice compared to WT (Rab27b^+/+^) littermate controls (Fig. 4c). LAL activity was similarly reduced in Rab27b KO neurons compared to WT neurons (Fig. 4d). Lysosomal numbers as determined by Lysotracker-positive counts were not different between WT and Rab27b neurons (Fig. 4e). To validate that these findings were driven by lysosomal activity and not confounded by differences in the uptake of quenched substrates, we assessed bulk endocytosis in Rab27b KO and WT neurons using 488-tagged dextran (10kDa). Dextran uptake was comparable between WT and Rab27b KO neurons (Fig. 4f).

We also examined whether increased αsyn load in primary neurons would exacerbate the lysosomal defects in primary hippocampal cultures. We crossed Rab27b KO mice with the transgenic A53T human αsyn mouse which demonstrates an approximately six-fold increase in αsyn levels compared to endogenous levels [36] (Fig. 5a). Lysosomal proteolytic activity as measured by DQ-BSA cleavage was reduced in Rab27b^-/-^ A53T negative (Rab27b KO/A53T^-^) and Rab27b^-/-^ A53T hemizygous (Rab27b KO/A53T^+^) neurons compared to either Rab27b^+/+^ A53T negative (WT) or Rab27b^+/+^ A53T hemizygous (A53T^+^) neurons, yet there was no additional impairment of DQ-BSA cleavage in Rab27b KO/A53T^+^ neurons compared to Rab27b KO neurons (Fig. 5b). Similarly, LAL activity was significantly reduced in both Rab27b KO and Rab27b KO/A53T^+^ neurons, but there was no difference between Rab27b KO and Rab27b KO/A53T^+^ neurons (Fig. 5c).

**Figure 5.**
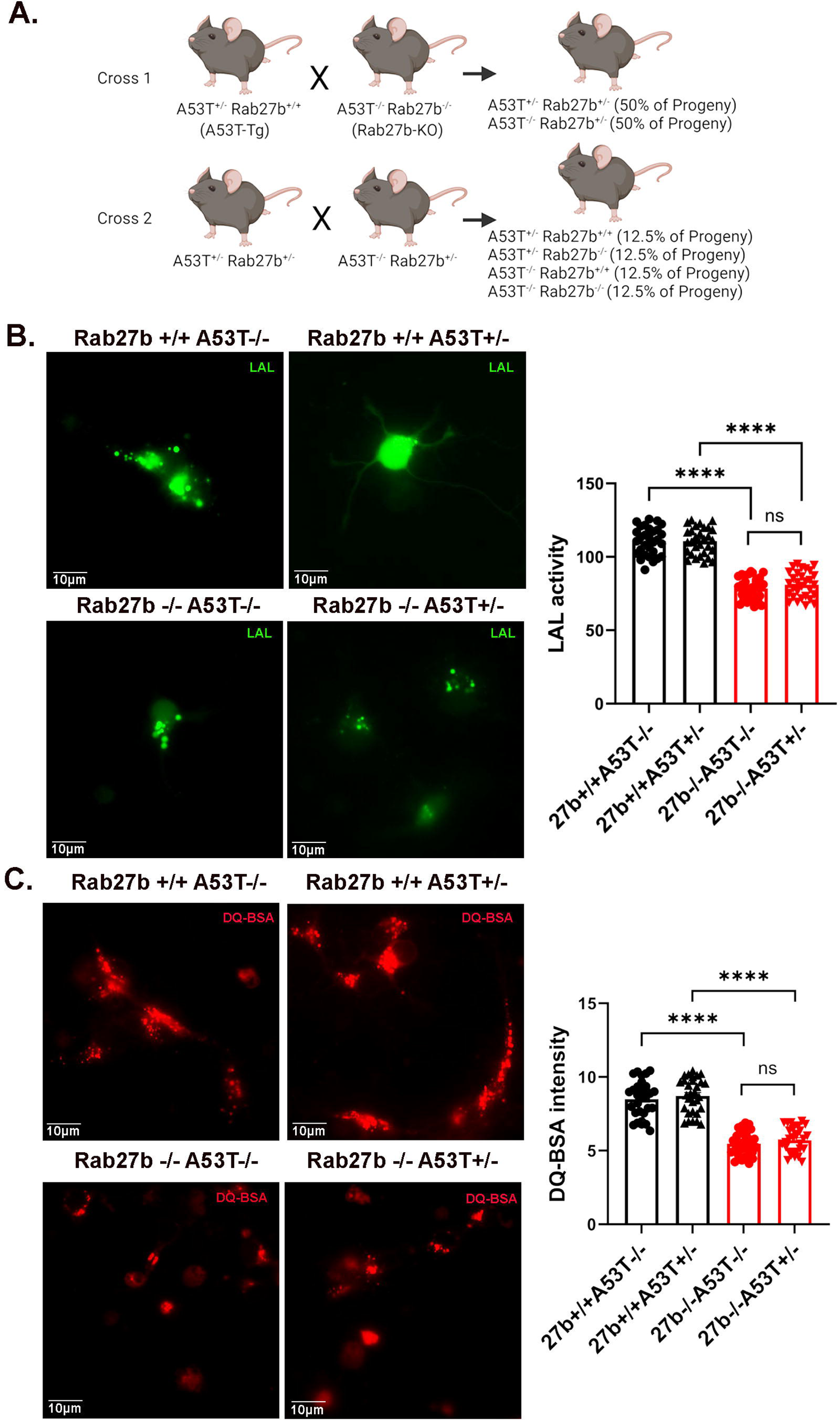
Rab27b KO x A53T hemizygous mice do not display exacerbated lysosomal pathology. A: Schematic depicting Rab27b KO x A53T hemizygous breeding strategy. B: Representative images and quantification of LAL activity in WT, A53T hemizygous, Rab27b KO, and Rab27b KO x A53T hemizygous neurons. n=30 individual cells across 3 independent rounds. One-way ANOVA with Tukey’s multiple comparison test; ****p<0.0001. C: Representative images and quantification of DQ-BSA activity in WT, A53T hemizygous, Rab27b KO, and Rab27b KO x A53T hemizygous neurons. n=30 individual cells across 3 independent rounds. One-way ANOVA with Tukey’s multiple comparison test; ****p<0.0001. All error bars denote SEM.

To examine if lysosomal defects were also present *in vivo*, we measured Ctsd and Gcase activity in cortical and hippocampal brain lysates from 2-4-month-old WT and Rab27b KO mice. Ctsd activity was reduced in both the cortex and hippocampus of Rab27b KO mice compared to WT littermates (Fig. 6a). Similarly, Gcase activity was reduced in the cortex and hippocampus of Rab27b KO mice compared to WT mice (Fig. 6b). Total mature Ctsd and Gcase protein levels were not changed (Fig. 6c-d), and we did not see any differences in LAMP-1 protein levels in WT vs Rab27b KO brain lysates (Fig. 6e-f).

**Figure 6.**
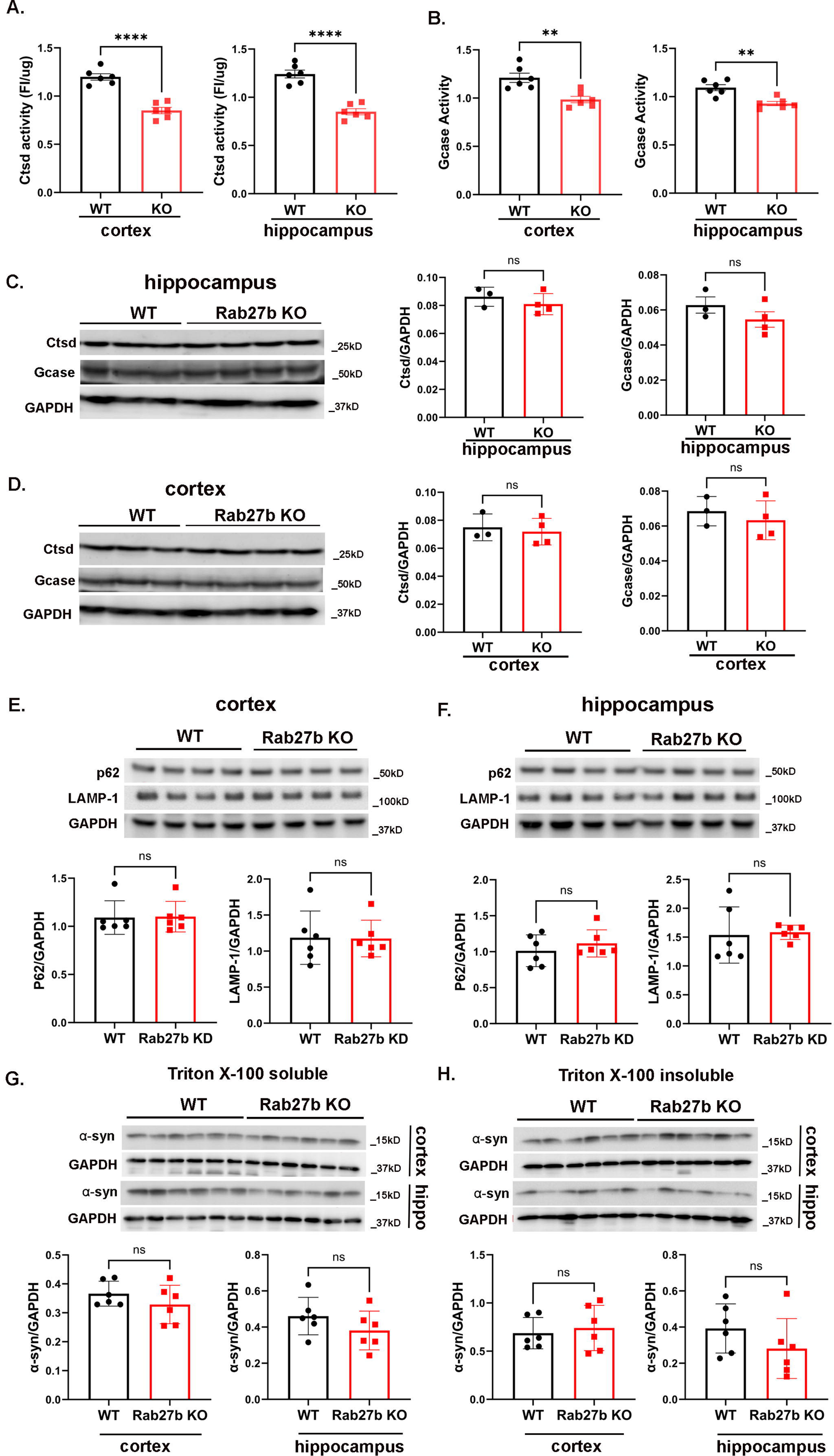
Rab27b KO mice show reduced lysosomal enzyme activity but no changes in autophagic-lysosomal markers or αsyn solubility. A: Quantification of Ctsd activity in cortical and hippocampal lysates from 2-4 month old WT and Rab27b KO mice. n=6 mice per group. Student’s t-test; ****p<0.0001. B: Quantification of Gcase activity in cortical and hippocampal lysates from 2-4 month old WT and Rab27b KO mice. n=6 mice per group. Student’s t-test; **p<0.01. C-D: Representative Western blots and quantification showing no change in mature Ctsd or Gcase expression levels in 2-4 month old Rab27 KO mice relative to WT controls. n=3-4 mice per group. Student’s t-test. E-F: Representative Western blots and quantifications of P62 and LAMP-1 expression in cortical and hippocampal lysates from 12-15 month old WT and Rab27b KO mice. n=6 mice per group. Student’s t-test. G-H: Representative Western blots and quantifications of αsyn levels in Triton X-100 soluble and insoluble cortical and hippocampal lysates from 12-15 month old WT and Rab27b KO mice. n=6 mice per group. Student’s t-test. All error bars denote SEM.

### Rab27b KO exacerbates αsyn pathology

We next examined whether Rab27b KO was sufficient to cause increases in insoluble αsyn in brain tissue. We measured Triton X-100-soluble and insoluble fractions from the hippocampus and cortex of Rab27b KO and WT littermates at 2-4 and 12-15 months of age. αSyn levels in both the Triton X-100 soluble and insoluble fractions were comparable in WT and Rab27b KO lysates at both ages (Fig. 6g-h). We also assessed the autophagic substrate marker p62 and found no changes in its expression in 12-15 month old mice (Fig. 6e-f).

We then examined Rab27b KO in the context of αsyn stress by assessing its effect on αsyn aggregation induced by αsyn preformed fibrils (PFFs) in culture. Primary hippocampal neurons from WT and Rab27b KO littermates were treated with αsyn PFFs at 1µg/ml at DIV 5, and then cultures were fixed and stained for phosphorylated αsyn at S129 (psyn) as a measure of aggregated αsyn as previously described [37]. We found that Rab27b KO primary neurons accumulated more psyn at 7 and 10 days post-PFF treatment (Fig. 7), indicating that KO of Rab27b can exacerbate αsyn-induced pathology.

**Figure 7.**
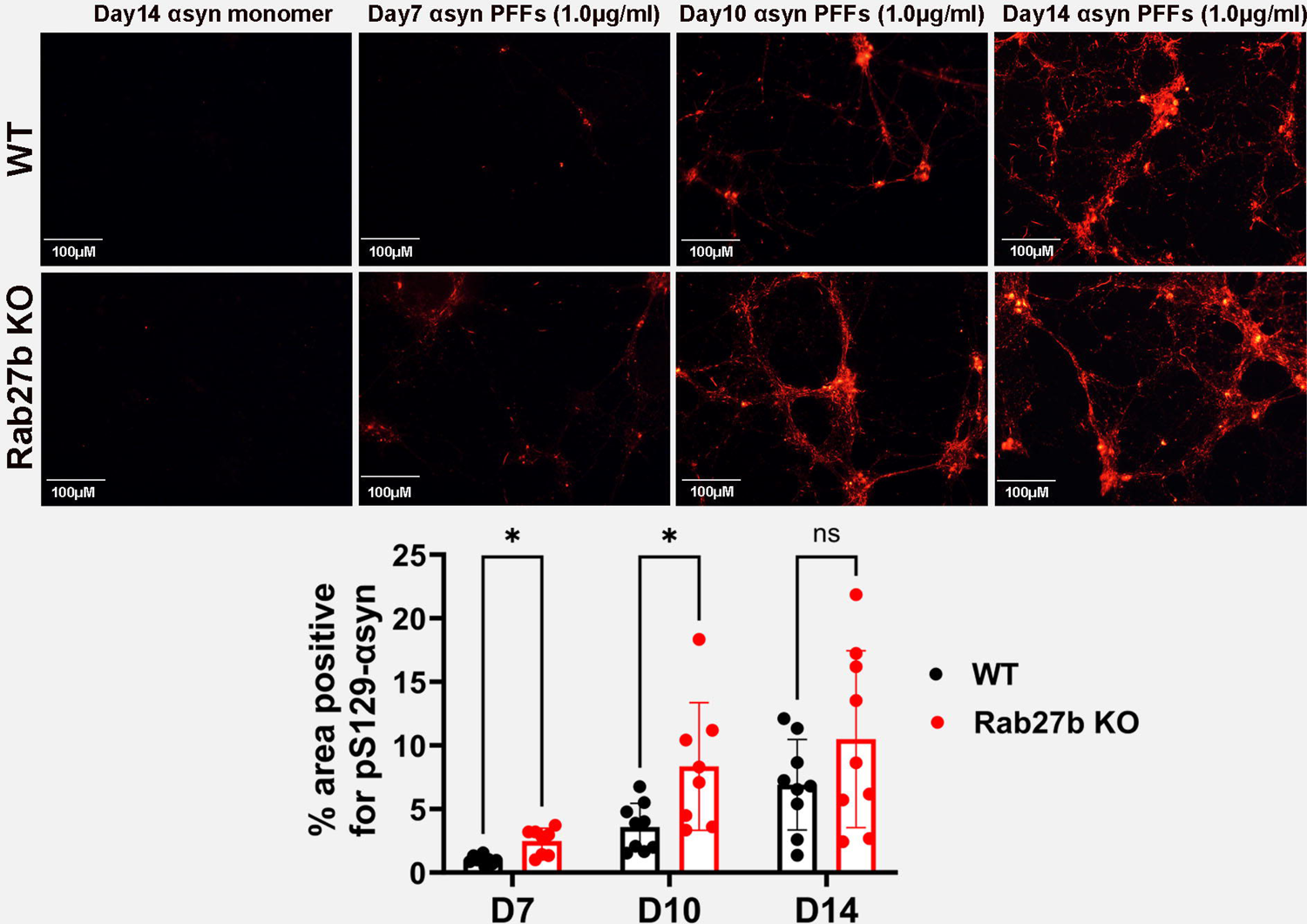
Rab27b KO causes increased accumulation of pS129 αsyn upon PFF treatment. Representative images and quantification showing pS129 immunostaining of WT and Rab27b KO hippocampal primary neurons over time after treatment with either monomeric or fibrillar αsyn. Unpaired t-tests with Holms-Sidak correction for multiple comparisons; *p<0.05.

### Rab27b KO impairs anterograde lysosomal trafficking in axons

Rab27b has been implicated in vesicular trafficking in axons [38]. To test if Rab27b KO could affect trafficking of acidic vesicles labeled with Lysotracker in axons, we performed live imaging of WT and Rab27b KO hippocampal neurons at DIV 8 (Fig. 8a-b). We found that the percentage and the density of cargo moving in the anterograde direction was significantly decreased in Rab27b KO neurons compared to WT (Fig. 8c-d). Additionally, the percentage of Lysotracker-positive vesicles that were stationary was increased in Rab27b KO neurons compared to WT (Fig. 8c). We also found that the net velocity of acidic vesicles was significantly reduced in both anterograde and retrograde directions in Rab27b KO neurons compared to WT (Fig. 8e). Although there was an increase in stationary tracks, the pause duration of Lysotracker-positive vesicles was not affected (Fig. 8f).

**Figure 8.**
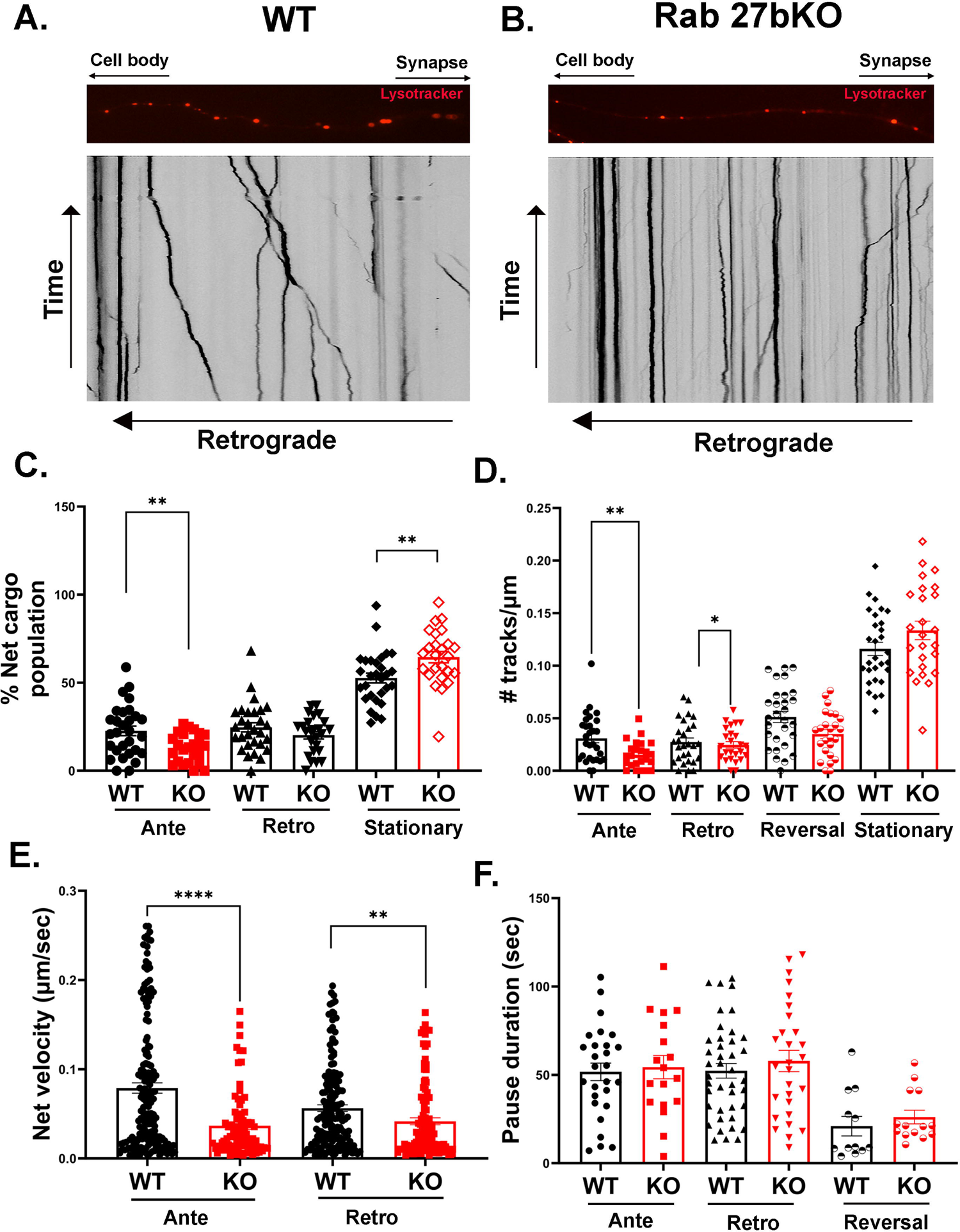
Rab27b KO impairs axonal acidic vesicle trafficking in primary neurons. A-B: Representative images and kymographs depicting the movements of Lysotracker-positive puncta in WT (A) and Rab27 KO (B) primary neurons. C-F: Quantification of percent net cargo population, number of tracks per µm (density), net velocity, and pause duration of Lysotracker-positive vesicles in WT and Rab27 KO primary neurons. Points represent individual tracks from six neurons per genotype measured across four independent rounds. Independent Student’s t-tests (C), Mann-Whitney U tests (E), or both (D, F) within each direction; *p<0.05, **p<0.01, ****p<0.001. All error bars denote SEM.

### Rab27b overexpression enhances lysosomal enzymatic function

We next determined if overexpression (OE) of Rab27b could enhance lysosomal activity. Isyn cells were transduced with either a GFP lentivirus or GFP-tagged Rab27b lentivirus which were both selectable by puromycin. We confirmed increased Rab27b expression in isyn cells transduced with GFP-tagged Rab27b lentivirus compared to GFP lentivirus (Fig. 9a). DQ-BSA cleavage was increased by 23% in isyn/Rab27b OE cells compared to isyn/GFP cells (Fig. 9b). Similarly, Ctsd and Gcase enzyme activities were increased by 36% and 15%, respectively, in isyn/Rab27b OE cells compared to isyn/GFP cells (Fig. 9c-d). We did not observe any changes in LAMP-1-positive vesicles or in the expression levels of Gcase or mature Ctsd between isyn/Rab27b OE cells and isyn/GFP cells (Fig. 9e-f).

**Figure 9.**
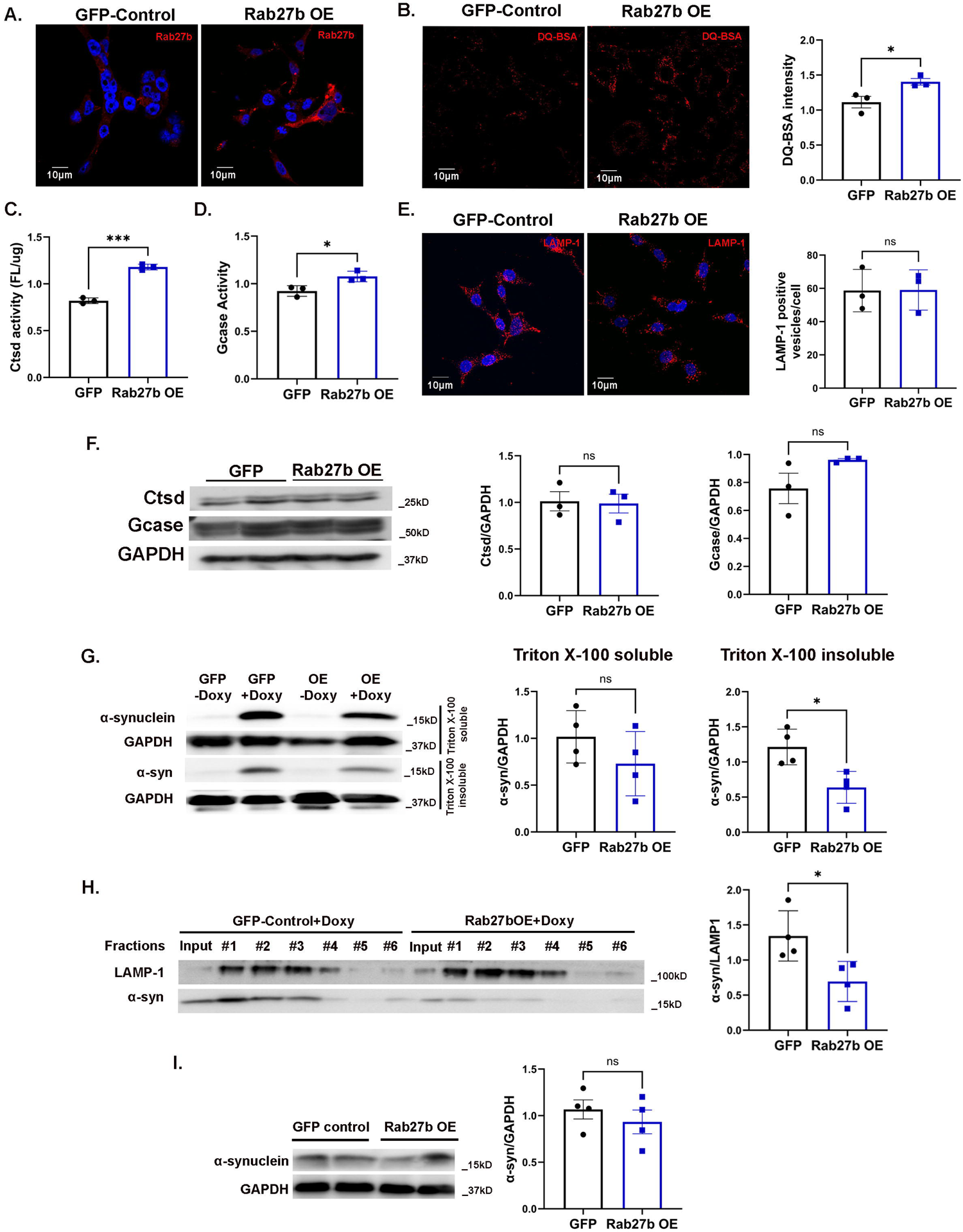
Rab27b OE cells display reduced αsyn pathology and enhanced lysosomal activity. A: Representative image showing immunostaining for Rab27b in isyn/GFP control and isyn/Rab27b OE cells. B: Representative images and quantification of DQ-BSA activity in isyn/GFP control and isyn/Rab27b OE cells. n=3 independent rounds. Student’s t-test; *p<0.05. C: Quantification of Ctsd activity in isyn/GFP control and isyn/Rab27b OE cells. n=3 independent rounds. Student’s t-test; ***p<0.001. D: Quantification of Gcase activity in isyn/GFP control and isyn/Rab27b OE cells. n=3 independent rounds. Student’s t-test; *p<0.05. E: Representative images and quantification of LAMP-1-positive puncta in isyn/GFP control and isyn/Rab27b OE cells. n=3 independent rounds. Student’s t-test. F: Representative Western blots and quantification of mature Ctsd and Gcase expression levels in isyn/GFP control and isyn/Rab27b OE cells. n=3 independent rounds. Student’s t-test. G: Representative Western blot and quantification of αsyn levels in Triton X-100 soluble and insoluble cell lysates from isyn/GFP control and isyn/Rab27b OE cells. n=4 independent rounds. Student’s t-test; *p<0.05. H: Representative Western blot and quantification of αsyn levels in LAMP-1-enriched fractions of isyn/GFP control and isyn/Rab27b OE cells. Fractions 1-6 denote successive fractions. αSyn levels were normalized to LAMP-1 enrichment. n=4 independent rounds. Student’s t-test; *p<0.05. I: Representative Western blot and quantification of baseline levels of αsyn in input controls of induced isyn/Rab27b OE and isyn/GFP control cells. Student’s t-test. All error bars denote SEM.

### Rab27b overexpression is protective against αsyn pathology

We previously determined that Rab27b KD in isyn cells results in increased intracellular Triton X-100-insoluble αsyn [32]. To determine if Rab27b OE could reduce insoluble αsyn accumulation, we prepared Triton-soluble and insoluble protein fractions from isyn/Rab27b OE and isyn/GFP cells. αSyn levels in the Triton-soluble fractions were not significantly changed between isyn/Rab27b OE and isyn/GFP cells, but Rab27b overexpression significantly reduced Triton-insoluble αsyn levels by 65% compared to control isyn/GFP cells (Fig. 9g).

Given the reduction in insoluble αsyn with Rab27b OE, we next assessed the levels of αsyn retained in lysosomal fractions. We found that isyn/Rab27b OE cells show reduced αsyn in lysosomally-enriched fractions compared to isyn/GFP cells, potentially indicating enhanced αsyn lysosomal degradation with Rab27b OE (Fig. 9h). Of note, total cellular lysate levels of αsyn in isyn/GFP vs isyn/Rab27b OE cells did not differ (Fig. 9i). Overall, isyn cells with Rab27b OE display enhanced lysosomal activity and reduced αsyn accumulation, indicating a potentially protective role of Rab27b against αsyn pathology.

### Rab27b is elevated in human Incidental Lewy Body Disease

We have previously shown that Rab27b is elevated in temporal cortical lysates from human DLB subjects relative to age-matched controls [32]. We evaluated whether Rab27b protein levels were altered at pre-clinical stages by measuring Rab27b levels in a cohort of Incidental Lewy Body Disease (iLBD) subjects. iLBD is viewed to represent the earliest stages of synucleinopathy and is pathologically defined by the presence of Lewy bodies or neurites in autopsied subjects without clinical diagnosis of Parkinsonism or dementia [39–42]. We found a 55% increase in Rab27b levels in temporal cortical lysates from iLBD patients relative to age-matched controls (Fig. 10). Based on these findings, we conclude that the increase in Rab27b levels we have previously observed in DLB and PD subjects is likely a compensatory mechanism that occurs early in disease.

**Figure 10.**
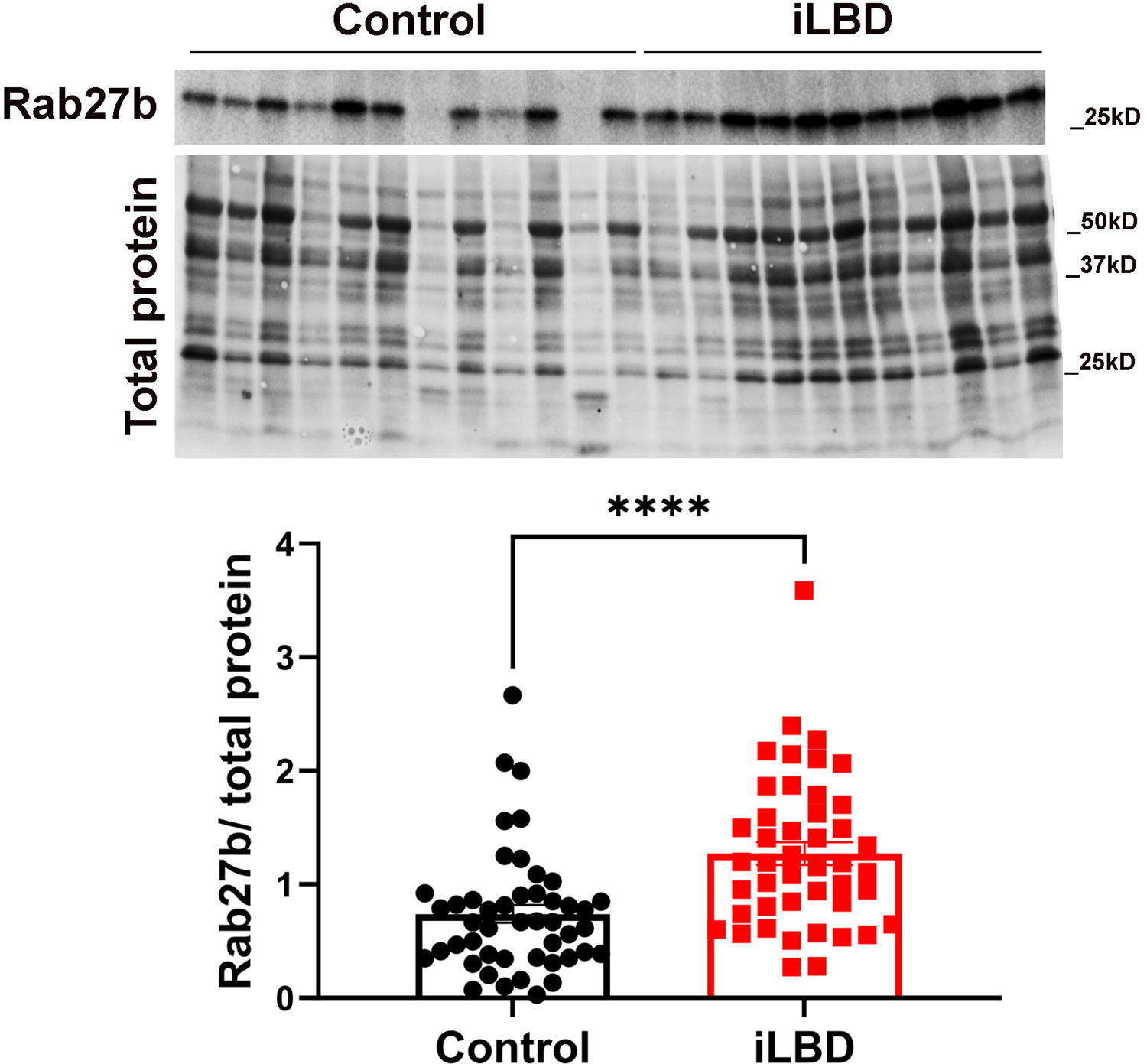
Rab27b is elevated in human incidental lewy body disease (iLBD). Representative western blot showing Rab27b levels and corresponding total protein per lane. Quantification of Rab27b levels in human temporal cortical lysates from iLBD subjects relative to age-matched controls. n=45. Student’s t-test; ****p<0.0001. Error bars depict SEM.

## DISCUSSION

Our findings point to a critical role for Rab27b in lysosomal function and the clearance of αsyn. We had previously shown that Rab27b promoted αsyn aggregation and toxicity using a doxy-inducible αsyn paracrine cell model. Here we show that Rab27b KD disrupted lysosomal function, which resulted in diminished αsyn clearance as demonstrated by increased αsyn levels in lysosomes. Similarly, we found lysosomal defects in primary neuronal cultures and in the cortex and hippocampus of Rab27b KO mice. Rab27b KO neurons showed increased αsyn pathology upon αsyn PFF treatment compared to WT neurons. We also observed increased Rab27b protein levels in temporal cortical lysates from subjects with iLBD. This suggests that increases in Rab27b may reflect compensatory mechanisms to promote lysosomal function in the brain in order to handle αsyn pathological stress. Indeed, we found that Rab27b OE enhanced lysosomal function in inducible αsyn cells and reduced levels of insoluble αsyn.

How Rab27b specifically enhances lysosomal function in neurons is still not fully clear, but our data points to alterations in vesicular transport. We found that Rab27b KD/KO did not alter lysosomal numbers, lysosomal pH, or lysosomal enzyme levels. We did find that Rab27b KO neurons showed disruptions in the anterograde transport of Lysotracker-positive vesicles within axons, suggesting that Rab27b is involved in trafficking of lysosomes to appropriate locations within the cell. A role for Rab27b in axonal transport is not entirely novel. For example, Rab27b has been implicated in the axonal transport of TrkB-positive vesicles [38]. A role for Rab27b in axonal transport could have significant impact on αsyn clearance. Effective protein clearance is dependent on bidirectional transport of autophagic vesicles and lysosomes in neurons. αSyn protein levels are enhanced at synapses, and evidence points to the initiation of αsyn pathology at the axon terminal [43–46], where degradative capacity is limited [47–49]. Protein clearance at distal terminals is dependent on the retrograde transport of autophagosomes that engulf misfolded proteins to the soma where they fuse with mature lysosomes [48–50]. Protein clearance is also dependent on the delivery of lysosomes to distal axons where local fusion with autophagic vesicles to initiate clearance occurs [47, 49]. Indeed, disruption of vesicular transport within axons has been shown to reduce αsyn clearance [47].

Another possible mechanism by which Rab27b could promote lysosomal function is that it may support the transport of mannose-6-phosphate receptor (M6PR)-positive vesicles containing immature lysosomal enzymes from the trans-Golgi network to lysosomes where they are cleaved and activated. We did not observe any measurable differences in expression levels or maturation state of Ctsd or Gcase, yet we were measuring total cellular enzyme levels. It is possible differences in enzymatic maturation states are present within lysosomes. Further work is needed to look specifically at enzyme maturation within lysosomes. Additionally, while Lysotracker labels lysosomes, it can also label other acidic vesicles, including late endosomes and autophagosomes. Future work will investigate what type of acidic vesicles are specifically affected by Rab27b KO and will also test whether axonal transport of other types of vesicles, such as M6PR-positive vesicles, are also affected by Rab27b.

Our finding that lysosomal function was diminished in Rab27b KO neurons even in the absence of excess αsyn suggests that an endogenous function for Rab27b in neurons is maintenance of lysosomal activity. This finding suggests that Rab27b could play a role in other neurodegenerative disorders marked by protein aggregation, such as Alzheimer’s disease, Huntington’s disease, and others. Evaluation of the impact of Rab27b is important to assess in other neurodegenerative disorders.

We previously found that Rab27b protein levels are increased in human PD and DLB brains. Here we provide additional data showing that Rab27b protein levels are also increased in iLBD brains. iLBD is viewed as a pre-clinical form of PD and DLB in which patients have evidence of Lewy Body pathology yet have not developed clear clinical signs of synucleinopathy. This finding that Rab27b is elevated at this pre-clinical stage suggests that increases in Rab27b reflect compensatory mechanisms by which neurons are trying to handle excess proteostatic stress from αsyn aggregates. Further boosting of Rab27b function could provide a therapeutic target for PD and DLB. This may be most practical in GBA-related disorders, in which Gcase activity is diminished and thought to be responsible for the increased risk in synucleinopathies. Given our findings that Rab27b OE can boost Gcase activity, mechanisms targeting Rab27b could be a therapeutic intervention for GBA.

In conclusion, we found that loss of Rab27b reduces lysosomal function in neuronal cell lines, primary neurons, and in brain tissue. Rab27b OE increases lysosomal function and reduces αsyn accumulation, pointing to Rab27b as a target for therapeutic intervention.

## METHODS

### Cell Lines and Mouse Models

#### Cell Lines

We previously developed a doxycycline (doxy)-inducible SK-N-BE(2)-M17 cell line (isyn) that overexpresses and secretes αsyn into the culture media upon treatment with doxycycline, as previously described [37]. To reduce Rab27b expression, isyn cells were transduced with a lentiviral Rab27b-targeted shRNA (5′-CCCAAATTCATCACTACAGTA-3′) to knock down Rab27b or with a lentiviral nontarget (nt) shRNA (Sigma-Aldrich SHC016), as a control as previously described [32].

To test the effect of Rab27b overexpression (OE), isyn cells were transduced with a lentiviral Rab27b-GFP construct (Genecopoeia EX-Q0376-Lv103) to overexpress Rab27b or a lentiviral GFP-expressing construct (Genecopoeia EX-NEG-Lv103) as a control. Cells were selected for stable transfection with puromycin (1µg/ml).

All isyn-based cell lines were treated with 500µg/ml G418 sulfate to maintain selection for inducible αsyn constructs, and with 1µg/ml puromycin to maintain selection for nt shRNA, Rab27b KD shRNA, GFP, and Rab27b OE constructs. Cells were maintained in a 1:1 ratio of F-12K and EMEM supplemented with 10% Fetal Bovine Serum and 1% penicillin-streptomycin. To induce αsyn overexpression, cells were treated with 10µg/ml doxycycline hyclate unless otherwise specified [51].

#### Rab27b KO Mice

Mice were used and maintained in accordance with University of Alabama at Birmingham (UAB) Institutional Animal Care and Use Committee (IACUC) protocols and the guidelines of the National Institute of Health (NIH). Rab27a and -b double knockout (DKO) mice acquired from Miguel Seabra [34] were crossed with C57BL/6-J mice from Jackson Laboratories (000664) to generate a single KO mouse for Rab27b. Rab27 DKO mice were crossed with WT mice to generate Rab27a^+/-^ Rab27b^+/-^ progeny, which were then crossed to each other to generate Rab27a^+/+^ Rab27b^+/-^ mice. For maintenance breeding, Rab27b^+/-^ heterozygous mice were bred together to generate litters with both WT control (Rab27b^+/+^) and Rab27b-KO (Rab27b^-/-^) pups for experimental use. Mice were genotyped for Rab27b and αsyn Exon 4 as previously described [35].

#### Rab27b KO A53T Transgenic Mice

Rab27b KO (Rab27b^-/-^) mice were crossed with A53T hemizygous (A53T^+/-^) mice [36] acquired from Jackson Laboratories (006823) to generate progeny that were Rab27b heterozygous (Rab27b^+/-^) and either A53T hemizygous (A53T^+/-^) or A53T negative (A53T^-/-^). These progeny were then crossed to generate the following gene combinations: Rab27b^+/+^ A53T^-/-^, Rab27b^+/+^ A53T^+/-^, Rab27b^-/-^ A53T^-/-^, and Rab27b^-/-^ A53T^+/-^. These animals were used to set up final crosses in two parallel lines: a Rab27b WT line (Rab27b^+/+^ A53T^+/-^ x Rab27b^+/+^ A53T^-/-^) and a Rab27b KO line (Rab27b^-/-^ A53T^+/-^ x Rab27b^-/-^ A53T^-/-^). Pups from these parallel cages were cultured together when born on the same day for experiments involving all four final genotypes: Rab27^+/+^ A53T^-/-^ (WT mice); Rab27b^+/+^ A53T^+/-^ (A53T^+^ mice); Rab27b^-/-^ A53T^-/-^ (Rab27b KO mice); and Rab27b^-/-^ A53T^+/-^ (Rab27b KO/A53T^+^ mice).

#### Primary neuronal cultures

Primary neurons were harvested from P0 pups as described previously [37]. Briefly, P0 pups were anesthetized on ice before brain collection. Hippocampi were bilaterally isolated and digested with papain for 30 minutes at 37°C. Hippocampi were mechanically dissociated and cells were plated in Neurobasal A media supplemented with B27, GlutaMax, Fetal Bovine Serum (FBS), and penicillin-streptomycin. At 24h, cells were treated with 1µm AraC in media without FBS to inhibit glial proliferation.

#### Immunocytochemistry

Isyn cells were plated on glass coverslips in 24-well plates at 10,000 cells per well and treated with 10µg/ml doxy for 96 hours before being harvested by fixation in a 4% paraformaldehyde (PFA) / 4% sucrose solution. Following washing, cells were permeabilized in 0.1% Triton X-100 in 3% BSA for 20 minutes followed by a one hour block in 3% BSA. Cells were incubated in primary antibodies (Rab27b, LAMP-1, LC3b, P62) in 3% BSA overnight at 4°C. Primary antibodies are described in Table 2. Cells were incubated with goat-anti-mouse or goat-anti-rabbit secondary antibodies in 3% BSA for one hour before being washed and mounted on slides with Prolong Diamond DAPI mounting media for imaging on a confocal microscope at x63 (Nikon Eclipse Ti2 scanning confocal microscope). Pearson’s correlation coefficients were quantified using Nikon NIS-Elements.

**Table 2.** Primary antibodies used.

| Primary Antibody | Dilution | Source | Identifier |
| --- | --- | --- | --- |
| LAMP-1 - Human | WB: 1:250<br>ICC: 1:100 | Iowa Developmental Studies Hybridoma Bank | H4A3 |
| LAMP-1 - Mouse | WB: 1:250<br>ICC: 1:100 | Iowa Developmental Studies Hybridoma Bank | 1D4B |
| αsyn | WB: 1:1000<br>ICC: 1:500 | BD Biosciences | 610787 |
| GAPDH | WB: 1:2000 | Cell Signaling | 14C10 |
| Rab27b | WB: 1:500<br>ICC: 1:500 | Proteintech | 13412-1-AP |
| Rab27b | PLA: 1:100 | abcam | ab-76779 |
| αsyn | PLA: 1:200 | Cell signaling | CS2642S |
| P62 | WB: 1:1000<br>ICC: 1:200 | Abcam | Ab56416 |
| LC3b | WB: 1:500<br>ICC: 1:200 | Sigma | L7543 |
| pS129 αsyn | ICC: 1:1000 | Abcam | Ab51253 |
| Ctsd | WB: 1:1000 | Invitrogen | MA1-26773 |
| Gba1 | WB: 1:1000 | Sigma | G4171 |

#### Proximity Ligation Assay (PLA)

Isyn cells were plated on glass coverslips in 24-well plates at 10,000 cells per well and treated with 10µg/ml doxy for 96 hours before being harvested by fixation in 4% PFA for 20 minutes at room temperature. PLA was executed according to the manufacturer’s protocols. Briefly, cells were incubated in primary antibody at 4°C overnight (see Table 2), then washed and treated with DuoLink PLA PLUS and MINUS probes before amplification and coverslipping. Cells were imaged on a confocal microscope at x63 as described above.

#### Lysosomal Fractionation: isyn/Rab27b KD cells

Isyn/Rab27b KD or control cells were treated with 10µg/ml doxy for 96 hours before being harvested for lysosomal fractionation using a Percoll density gradient as previously described [52]. Briefly, cells were manually lysed and added to a sucrose-Percoll gradient before being spun at 36,000*xg* for 30 minutes (Beckmann-Coulter Optima XE-90 Ultracentrifuge; SW 31 Ti rotor). Lysosomal fractions were manually collected and used for Western blot analysis. Blots were probed for LAMP-1 and αsyn (Table 2).

#### Lysosomal Fractionation: isyn/Rab27b OE cells

Isyn/Rab27b OE or isyn/GFP control cells were treated with 10µg/ml doxy for 96 hours before being harvested for lysosomal fractionation using a Lysosome Enrichment Kit (Table 3). Briefly, cells were lysed and added to an OptiPrep density gradient before being spun at 145,000*xg* for two hours (Beckmann-Coulter Optima XE-90 Ultracentrifuge; SW 31 Ti rotor). Lysosomal fractions were manually collected, washed, and then further lysed to free lysosomal proteins before Western blot analysis. Blots were probed for LAMP-1 and αsyn (Table 2).

**Table 3.** Other reagents used.

| **Reagent** | **Source** | **Identifier** |
| --- | --- | --- |
| **Puromycin** | Fisher Scientific | BP2956100 |
| **G418 Sulfate** | KSE Scientific | 108321-42-2 |
| **F-12K Media** | ATCC | 30-2004 |
| **EMEM Media** | ATCC | 30-2003 |
| **Fetal Bovine Serum** | Sigma | F2442 |
| **Penicilin-Streptomycin** | Cytiva | SV30010 |
| **Doxycycline hyclate** | Sigma | D9891 |
| **Papain** | Worthington Biochemical | #3176 |
| **Neurobasal A Media** | Gibco | 21-203-049 |
| **B27** | Gibco | 12587001 |
| **GlutaMax** | Gibco | A1286001 |
| **AraC** | Cytarabine; LGC Standards | EPC3350000 |
| **Lysosome Enrichment Kit** | Thermo Scientific | 89839 |
| **RIPA lysis buffer** | Thermo Scientific | 89901 |
| **Ctsd Substrate** | Sigma | 219360 |
| **Pepstatin A** | Sigma | 77170 |
| **35mm Mat-Tek dishes** | Mat-Tek | P35G-0-14-C |
| **LysoLive LAL Kit** | abcam | 253380 |
| **DQ-BSA Substrate** | Invitrogen | D12051 |
| **Neurobasal A - Phenol Red media** | Gibco | 12349015 |
| **Dextran 10,000 MW** | Invitrogen | D22910 |
| **Trypan Blue** | Sigma | T6146 |
| **BlueSafe gel stain** | Thermo Scientific | 24594 |
| **4-MU β-D-glucopyranoside** | Sigma-Aldrich | M3633 |
| **Conduritol B Epoxide** | Sigma-Aldrich | C5424 |
| **Prolong Diamond + DAPI** | ThermoFisher Scientific | P36962 |
| **Lysosomal Acidic pH Detection Kit** | Dojindo Labs | L266-10 |
| **BlueSafe Gel Stain** | Thermo Scientific | 24594 |
| **Lysotracker Red** | Invitrogen | L7528 |
| **Duolink PLA Probe Anti-Rabbit PLUS** | Sigma-Aldrich | DUO92002 |
| **Duolink PLA Probe Anti-Mouse MINUS** | Sigma-Aldrich | DUO92004 |
| **Duolink In Situ Detection Reagents Red** | Sigma-Aldrich | DUO92008 |

### Lysosomal Enzyme Assays

#### Cathepsin D activity

Isyn-based cells were treated with 10µg/ml doxy for 96 hours before being lysed in RIPA buffer to extract proteins. 5µg of protein was incubated with either 40µM Ctsd Substrate or 40µM Ctsd Substrate plus 30µM Substrate Inhibitor Pepstatin A diluted in 50mM Sodium Acetate (pH 4.0) for 60 minutes at 37°C in 96 well plates. Plates were read on a SpectraMax iD3 (Molecular Devices) plate reader (EX: 360nm EM: 460nm), and substrate wells were normalized to substrate and inhibitor wells. Similarly, brain lysates from WT and Rab27b KO mice were incubated with Ctsd Substrate with and without Pepstatin A to measure Ctsd activity *in vivo*.

#### Glucocerebrosidase activity

Isyn-based cells were treated and lysed as described above. Gcase activity was measured as previously described [53]. Briefly, 5µl of each sample was added to a 96 well plate and incubated with 45µl of Gcase assay buffer (1.11% BSA, 0.28% Triton X-100, 0.28% Taurocholic Acid, 1.11 mM EDTA) containing either 1.11mM 4-methylumbelliferone β-D-glucopyranoside (4-MUGluc) or 1.11mM 4-MUGluc with 0.2mM Conduritol B Epoxide (CBE). Plates were incubated at 37°C for one hour, after which 50µl of stop solution (0.4M Glycine pH 10.8) was added to each well. Fluorescence intensity was recorded on a SpectraMax iD3 (Molecular Devices) plate reader (EX: 355nm EM: 460nm) and substrate wells were normalized to CBE inhibitor wells and then to total protein levels. Similarly, brain lysates from WT and Rab27b KO mice were incubated with 4-MUGluc Substrate with and without CBE to measure Gcase activity *in vivo*.

#### Lysosomal Acid Lipase activity

Isyn-based cells were plated on 35mm Mat-Tek live imaging dishes and treated with 10µg/ml doxy for 96 hours. Cells were then treated with LAL substrate for six hours before being imaged live on a Carl Zeiss Observer high-speed live cell imaging system microscope. For examination of LAL activity in primary hippocampal neurons, neurons were treated at DIV 8 with LAL substrate prior to imaging.

#### DQ-BSA assay

Isyn-based cells were plated on 35mm Mat-Tek live imaging dishes and treated with 10µg/ml doxy for 96 hours. Cells were then treated with DQ-BSA substrate and imaged after six hours using either a Carl Zeiss Observer high-speed live cell imaging system or Nikon Eclipse Ti2 scanning confocal microscope. WT and Rab27b KO primary hippocampal neurons were treated at DIV 8 prior to treatment and imaging.

#### Lysotracker staining

Isyn-based cells were plated on 35mm Mat-Tek live imaging dishes and treated with 10µg/ml doxy for 96 hours. Cells were then treated with 1µM Lysotracker Red and incubated for one hour. Then the cells were live imaged at x63 using a Carl Zeiss Observer high-speed live cell imaging system.

#### Lysosomal pH Assays

Isyn/Rab27b KD and control cells were plated on 35mm Mat-Tek live imaging dishes and treated with 10µg/ml doxy for 96 hours. Cells were then stained using a lysosomal pH kit from Dojindo labs according to the manufacturer’s protocols (Table 3). Cells were washed twice in serum-free media and then incubated in LysoPrime Green stain at 1:2000 for 30 minutes at 37°C to mark total lysosomes. Cells were then washed twice and incubated in pHLys Red stain at 1:1000 for 30 minutes at 37°C to mark acidified lysosomes. Cells were washed two more times, incubated with DAPI at 1:2000 for five minutes at 37°C, washed two more times, and then live-imaged in fresh media at x63 on a confocal microscope (Nikon Eclipse Ti2 scanning confocal microscope). Control cells incubated with BafA at 1:1000 were treated with LysoPrime Green as described above and then treated with a solution combining pHLysRed and BafA to inhibit lysosomal acidification before undergoing washes and imaging as described.

#### Triton X-100 fractionation

Isyn cells were treated with 10µg/ml doxy for 96 hours before being harvested. Control cells did not receive doxy. Cells were first resuspended and sonicated in lysis buffer (50mM TrisHCL pH 7.4, 175mM NaCl, and 5mM EDTA pH 8.0) with 1% Triton X-100 and spun for 60 minutes at 15,000*xg* to extract Triton X-100-soluble proteins. Pellets were then resuspended in lysis buffer containing 2% SDS and sonicated to extract Triton X-100-insoluble proteins. Mouse brains were homogenized and sonicated in lysis buffer with 1% Triton X-100, incubated on ice for 30 minutes, and spun for 60 minutes at 15,000*xg* to extract Triton X-100-soluble proteins. Pellets were then resuspended in lysis buffer containing 2% SDS and sonicated to extract Triton X-100-insoluble proteins. Samples were assessed for αsyn and GAPDH by Western blot (Table 2).

#### Dextran Internalization Assay

WT and Rab27b KO primary hippocampal neurons were plated on 35mm Mat-Tek dishes. At DIV 7, cells were placed on ice, washed twice in Neurobasal A - Phenol Red media and treated with 50mg/ml Dextran (10,000 MW). Cells were left on ice for 30 minutes to allow dextran to settle and were then moved to a 37° incubator for 30 or 60 minutes to allow endocytosis to occur. Plates were washed twice with Neurobasal A – Phenol Red, and extracellular signal was quenched with 1mM trypan blue for three minutes before cells were live-imaged on a Nikon Eclipse Ti2 scanning confocal microscope.

#### Lysosomal Trafficking Assay

WT and Rab27b KO primary hippocampal neurons were cultured on 35mm Mat-Tek dishes. At DIV 10, cells were exposed to 1µM Lysotracker Red for one hour. Axons from pyramidal neurons were identified based on their growth cone morphology, lack of spines, and non-tapering neurite structure. Axons were imaged using a ZEISS Z1 Cell Observer (63x oil, 1.4 NA) equipped with a Colibri2 cool LED 5-channel fast wavelength switching system and a Hammamatsu Orca Flash high-speed camera to assess axonal transport. Time lapse videos were acquired at 3 frames/sec x 2min. Videos of the axons were recorded for >100µm from the soma and from the axon terminal/synapses. Kymographs from time-lapse videos were generated using Kymoanalyzer [54]. This software provides parameters including the percentage of cargo in anterograde, retrograde, reversing, or stationary motion; the percentage of time spent in each type of motion; the number and frequency of pauses; the number and frequency of switches in direction; as well as net and segmental velocities.

#### Human Brain Lysates

Flash-frozen human temporal cortical brain samples from patients with Incidental Lewy Body Disease and age-matched controls were acquired from the Banner Sun Health Research Institute Brain and Body Donation Program. The Institutional Review Board (IRB) at UAB determined the use of human brains not human subjects research as tissue was from deceased persons. Samples were assessed for Rab27b and total protein (BlueSafe gel stain) via Western blot.

#### Pre-formed αsyn Fibril Assay

Pre-formed αsyn fibrils were prepared as previously described [37]. Before use, PFFs were diluted to 1mg/ml and sonicated using a QSonica700 cup horn sonicator with a 16°C water bath for 15 minutes at a 30% amplitude with cycling three seconds on and two seconds off pulses. Fibril fragmentation of fibrils was verified using dynamic light scattering before sonicated fibrils were added to primary neuronal cultures at 1µg/ml in neuronal media. Fibrils were added to hippocampal primary neurons at DIV-5, and cells were fixed in 4%PFA / 4%Sucrose / 1% Triton X-100 and stained with a pS129-αsyn antibody at seven, 10, and 14 days post treatment as previously described [37].

#### Western Blotting

Western blotting was performed as previously described [37].

#### Experimental Design and Statistical Analyses

GraphPad Prism 10 was used for statistical analysis of all experiments. All datasets were analyzed via Student’s t-test or one-way ANOVA with post-hoc Tukey’s multiple comparison testing post-hoc. *In vitro* PFF experiments were analyzed by multiple unpaired t-tests with correction for multiple comparisons using the Holm-Sidak method.

## ACKNOWLEDGEMENTS

We thank Drs. Geidy Serrano and Thomas Beach at the Banner Sun Health Research Institute Brain and Body Donation Program of Sun City, Arizona for the provision of human brain tissue. This study was supported by NIH [RF1NS115767 (TAY); R01 NS112203 (TAY); P50 NS108675 (TAY); T32 5T32GM135028 (KS); T32 5T32NS095775 (KS)] and the American Parkinson Disease Association.

## AUTHOR CONTRIBUTIONS

KS and RP designed and performed experiments, analyzed data, and wrote manuscript. ER and AN performed experiments and reviewed manuscript. FSP analyzed data and reviewed manuscript. TY designed experiments, analyzed data, and edited and finalized manuscript draft.

